# Identification of leptospiral protein antigens recognized by WC1^+^ *γδ* T cell subsets as target for development of recombinant vaccines

**DOI:** 10.1101/2021.07.16.452762

**Authors:** Aline F. Teixeira, Alexandria Gillespie, Alehegne Yirsaw, Emily Britton, Janice C. Telfer, Ana Lucia do Nascimento, Cynthia L. Baldwin

**Author notes:** Corresponding author, Avenida Vital Brazil, 1500, - 05503-900, Sao Paulo, SP, Brazil.

## Abstract

Pathogenic *Leptospira* species cause leptospirosis, a neglected zoonotic disease recognized as a global public health problem. It is also the cause of the most common cattle infection that results in major economic losses due to reproductive problems. γδ T cells play a role in the protective immune response in livestock species against *Leptospira* while human γδ T cells also respond to *Leptospira*. Thus, activation of γδ T cells has emerged as a potential component for optimization of vaccine strategies. Bovine γδ T cells proliferate and produce IFN-γ in response to vaccination with inactivated leptospires and this response is mediated by a specific subpopulation of the WC1-bearing γδ T cells. WC1 molecules are members of the group B scavenger receptor cysteine rich (SRCR) superfamily and are composed of multiple SRCR domains, of which particular extracellular domains act as ligands for *Leptospira*. Since WC1 molecules function as both pattern recognition receptors and γδ TCR coreceptors, the WC1 system has been proposed as a novel target to engage γδ T cells. Here, we demonstrate the involvement of leptospiral protein antigens in the activation of WC1^+^ γδ T cells and identified two leptospiral outer membrane proteins able to interact directly with them. Interestingly, we show that the protein-specific γδ T cell response is composed of WC1.1^+^ and WC1.2^+^ subsets, although a greater number of WC1.1^+^ *γδ* T cells respond. Identification of protein antigens will enhance our understanding of the role γδ T cells play in the leptospiral immune response and in recombinant vaccine development.

## Introduction

Leptospirosis is a neglected zoonotic disease recognized as a global public health problem that affects mainly rural farmers and urban slum dwellers. This disease is also considered one of the most common cattle infections and results in major economic losses due to reproductive problems such as abortions and stillbirths (Faine et al., 1999). The leptospirosis etiologic agent is the pathogenic spirochetes of the genus *Leptospira* (Bharti et al., 2003). Animals, especially rats, act as carriers, shedding leptospires through their urine into the environment. Both animals and human are infected from accidental direct contact or indirectly by exposure to the contaminated environment (Ko et al., 2009). After infection, individuals can exhibit a mild illness or develop severe complications due to involvement of multiple organ systems. Since leptospirosis’ initial symptoms are often undifferentiated from other febrile illnesses, early diagnosis may not occur and, consequently, result in ineffective treatment. The prevention of leptospirosis through vaccination is an alternative to post-infection treatment and can control the spread of disease. However, for the most part vaccines available for leptospirosis are based on whole cell killed or inactivated bacteria and the immune protection relies on antibody production against lipopolysaccharides (LPS) of the serovars present in the vaccine formulation. As LPS is unable to stimulate a classical T cell response, these vaccines promote a short-term immunity requiring annual boosting (Laurichesse et al., 2007; Martínez et al., 2004) (Yan et al., 2003; Yanagihara et al., 2007). The search for new vaccine strategies promoting a lasting immunological memory could contribute to control of leptospirosis.

Activation of nonconventional T cells, such as γδ T cells, are an alternative or supplement to conventional vaccines for infections that requires cell-mediated immune responses, since these cells can produce pro-inflammatory cytokines that may direct the αβ T cell response toward Th1 responses (Baldwin et al., 2014). It has been shown that both human and bovine γδ T cells are able to proliferate and produce IFN-γ in response to *Leptospira* (Brown et al., 2003; Klimpel et al., 2003; Naiman et al., 2001). In cattle, this response is mediated by a specific subpopulation of the WC1-bearing γδ T cells, which have a memory response (Blumerman et al., 2007). In some animals, this response was maintained for more than 2 years and *in vitro* stimulation of these cells promoted the expression of cell surface markers similar to those observed for the memory CD4 T cell population. Interestingly, γδ T cell responses were totally dependent on CD4 T cells, but a direct contact between cells was not required (Blumerman et al., 2007; Naiman et al., 2002). From these studies, it is evident that γδ T cells play an important role to leptospirosis infection and represent an interesting target in vaccine design. Moreover, the high frequency of this cell population in cattle makes it an excellent model for studying their involvement in responses to antigens from *Leptospira*.

Bovine WC1^+^ γδ T cells are CD2^-^, CD4^-^ and CD8^-^, and thus this γδ T cell subpopulation is characterized by the presence of the receptor WC1 (Baldwin et al., 2020; Telfer and Baldwin, 2015) WC1 is a transmembrane glycoprotein containing either 11 or 6 extracellular domains in cattle and expressed only on the surface of γδ T cells (Clevers et al., 1990; Herzig and Baldwin, 2009). WC1 belongs to the scavenger receptor cysteine rich (SRCR) superfamily and is closely related to SCART molecules found on murine γδ T cells and to CD163c-α in humans (Baldwin et al., 2014). It has been shown that members of the group B SRCR family interact directly with pathogens, including WC1 (Fabriek et al., 2005; Fabriek et al., 2009; Sarrias et al., 2007). The bovine WC1 gene family has 13 members classified into serologically-defined groups known as WC1.1 and WC1.2; cells in the WC1.1^+^ γδ T cell subpopulation largely comprise those that respond to *Leptospira* (Rogers et al., 2005a; Rogers et al., 2005b). The fundamental roles of WC1 molecules in response to *Leptospira* have been demonstrated by genetic silencing (Wang et al., 2011) and by their ability to act as pattern recognition receptors (PRR) with specific leptospira-binding domains. Only WC1 domains expressed by the *Leptospira*-responsive γδ T cell subset were able to bind to several species and serovars of *Leptospira* (Hsu et al., 2015). The TCR is also involved in *Leptospira* recognition, as shown by antibody blocking (Blumerman et al., 2006) and by antibody crosslinking of WC1 with the TCR-CD3 complex (Hanby-Flarida et al., 1996; Hsu et al., 2015) potentiating T cell activation. Thus, it is assumed that co-crosslinking of the γδ TCR and WC1 by *Leptospira* would be necessary for full activation of the WC1.1^+^ γδ T cells (Baldwin et al., 2020). However, the leptospiral components involved in this interaction are still unknown and discovery of these antigens is fundamental for development of recombinant vaccines that engage γδ T cells.

Many scavenger receptors bind to common pathogen-associated molecular patterns (PAMPs), such as lipopolysaccharides and lipoteichoic acids (Fabriek et al., 2009; Miró-Julià et al., 2011; Sarrias et al., 2007). In some cases, specific bacterial surface proteins are reported to be involved in these interactions (Loimaranta et al., 2009; Loimaranta et al., 2005). It seems the repertoire of antigens able to stimulate γδ T cells is also large, since other studies have shown γδ T cell interaction with the bacterial cell wall components, phosphoantigens, heat-shock proteins and protein complexes (Rhodes et al., 2001; Vesosky et al., 2004; Welsh et al., 2002) (Chen, 2013; O’Brien et al., 1992). However, how γδ T cells are activated by these antigens remains unknown. Interestingly, it was demonstrated for the first time that γδ T cells were able to respond directly to peptide antigens of *Mycobacterium bovis* and that this recognition required direct contact with APCs and recognition via the γδ TCR but that it occurs independently of antigen-presentation via MHC class I or II (McGill et al., 2014). This reinforces that paradigm that γδ TCR specificity and antigen interaction does not occur the same way as for αβ TCRs.

Generally, pathogens’ surface-exposed molecules are attractive targets for vaccines that rely on antibodies for efficacy due to the interactions of surface-exposed molecules with their host cells and tissues. The leptospiral outer membrane has many interesting characteristics. It is rich in LPS, unlike most other spirochetes. However, these molecules present unique structural features which differentiates them from other Gram-negative bacteria. As for other spirochetes, the leptospiral outer membrane is composed of many lipoproteins, although most of them have unknown functions. It has been suggested that bacterial lipoproteins can acts as PAMPs and be recognized by specific PRR (Takeda et al., 2002; Takeuchi et al., 2002). In this regard, we investigated the leptospiral antigens that could activate γδ T cell subsets and the role of WC1 in this activation. We demonstrated that γδ T cell from cattle immunized with an inactivated leptospiral vaccine proliferated and specifically responded to leptospiral protein antigens and, preferentially, WC1.1^+^ γδ T cells subsets were involved in this response. The discovery of these antigens will contribute to the design of vaccines against leptospirosis and understanding of the role that γδ T cell may play in these vaccine formulations.

## Material and Methods

### *Leptospira* strain

*Leptospira* reference serovar Copenhageni, strain M-20, in a semi-solid culture was received from the United States Department of Agriculture National Veterinary Services Laboratories (Ames, IA). It was cultivated from Dinger’s disk in 3 mL of Elinghausen-McCullough-Johnson-Harris (EMJH) medium (BD, Difco) supplemented with 10% Leptospira enrichment EMJH medium (BD, Difco) for 12 days at 30 °C. Following that, 3 mL of grown culture was added in 8 mL fresh EMJH medium until turbid. The propagation to larger volume was performed by a dilution of 1/10 and the culture was incubated for 7-12 days at 30 °C.

### Preparation of antigens from *L. interrogans*

To obtain the whole cell lysate (WCL) fractions a total of 100 mL of culture were centrifuged at 3,000 x g for 30 min at room temperature and the pellet washed with 10 mL PBS. The cell suspension was centrifuged again for 15 min and the cells were resuspended in 1 mL PBS buffer containing protease inhibitor (Halt Protease Inhibitor Cocktail 100x – ThermoFisher), DNase (Amresco) and RNase (Promega). Cells were lysed by three successive rounds of sonication for 30 sec at 20 kHz (Sonicator Ultrasonic processor XL 2020 – Heat systems) and centrifuged at 3,400 x g for 15 min at 4 °C to remove unbroken cells. The supernatant was collected and kept at −20 °C. Secreted protein fractions were also obtained from 100 mL culture. Cells were harvested by centrifugation as described above, the supernatant discarded, and the pellet washed with 10 mL PBS. Cells harvested by centrifugation were resuspended in 10 mL PBS 1x and kept at 30 °C for 4 h for secretion of residual proteins. The supernatant was recovered by centrifugation at 3,000 x g for 30 min at room temperature and the cells were resuspended again in 10 mL PBS 1 x and incubated overnight at 30 °C. Cell suspensions were centrifuged at 3,000 x g for 30 min at room temperature and the supernatant recovered. Both samples were concentrated 10x by using Amicon ultrafiltration with a molecular weight cut off 10 kDa (Millipore) and dialyzed in 10 mM ammonium bicarbonate. Preparations containing SDS-soluble cell wall proteins were performed as described to obtain the whole cell fraction. After centrifugation to remove unbroken cells, the supernatant collected was submitted to ultra-centrifugation at 27,000 x g for 20 min at 4 °C. The cell wall-containing pellet was resuspended in PBS with 2% SDS and stirred overnight at 37 °C. SDS soluble fractions were collected after centrifugation at 17,000 x g for 20 min at room temperature. The SDS extraction was repeated once for 4 h at 37 °C. SDS present on samples were removed by using Extract gel columns and the SDS-Out SDS precipitation kit (Thermo Scientific) (Hirschfield et al., 1990). Protein concentrations for each fraction were determined by the BCA protein assay kit (Thermo Scientific).

### Cloning, expression, and purification of recombinant proteins

Genomic DNA was extracted from *Leptospira interrogans* serovar Copenhageni strain M20 culture by using DNeasy Blood & Tissue kit (Qiagen) and used as a template to amplify *lipl21, lipl32, lipl41* and *lipl46* genes by PCR. Oligonucleotide primer pairs specific for each gene can be found in Table 1. The gene sequence was amplified without the signal sequence and then PCR amplicons were ligated into the *E.coli* expression vector pET28a (Novagen) at the restriction sites shown in Table 1. Positive clones were confirmed by commercial sequencing at GENEWIZ. Plasmids containing genes of interest were used to transform *E.coli* BL21 (DE3) Star (Invitrogen). Recombinant proteins were induced upon addition of 1mM IPTG in cultures containing kanamycin 50 *μ*g/mL and incubated with agitation at 37 °C. After 3 h induction, the cells were harvested by centrifugation, and the resulting bacterial pellet was resuspended in B-PER - Bacterial Protein Extraction reagent (ThermoFisher). Separation of soluble proteins from inclusion bodies was performed by centrifugation at 12,000 x g for 15 min at 4 °C and the insoluble fraction was resuspended in a PBS buffer containing 8M urea. The proteins were then purified by immobilized metal affinity chromatography by using HisPur Ni-NTA (Purify His-tagged recombinant fusion proteins using high-capacity nickel-nitrolotriacetic acid methal-chelate beads) spin columns (ThermoFisher). Recombinant proteins were refolded on the column by gradually removing urea until reaching to a concentration 0 M and then dialyzed against PBS. Protein concentrations were determined by NanoDrop A280. The efficiency of the expression and purification of recombinant proteins were evaluated by 12% SDS-PAGE and Western blotting by using anti-his tag antibodies (1:3,000) (Thermo Scientific) and HRP-conjugated goat anti-mouse IgG (1:10,000) (Southern Biotech), followed by ECL reagent kit chemiluminescence substrate (BioRad). For cloning individual WC1 domains, DNA constructs containing the WC1-3a1, WC1-3b2, WC1-3d6, WC1-3c8 and WC1-3e10 domains were used as template to amplify the genes by PCR (Chen et al., 2009; Hsu et al., 2015). Purified PCR products were ligated into the *E.coli* expression vector pETSUMO as described above for cloning into pET28a vector. Recombinant proteins were induced in SHuffle Express strains (New England Biolabs) upon addition of 1mM IPTG in cultures containing kanamycin 50 *μ*g/mL and incubated with agitation at 37 °C. After 3 h induction or overnight, the cells were harvested by centrifugation as described above. All proteins were purified by immobilized metal affinity chromatography from soluble fraction. Efficiency of the expression and purification of recombinant proteins were determined by 15% SDS-PAGE and Western blotting by using anti-his tag antibodies (1:10,000) (Sigma). Protein concentrations were quantified by comparison with a curve of BSA resolved by SDS-PAGE.

**Table 1.**
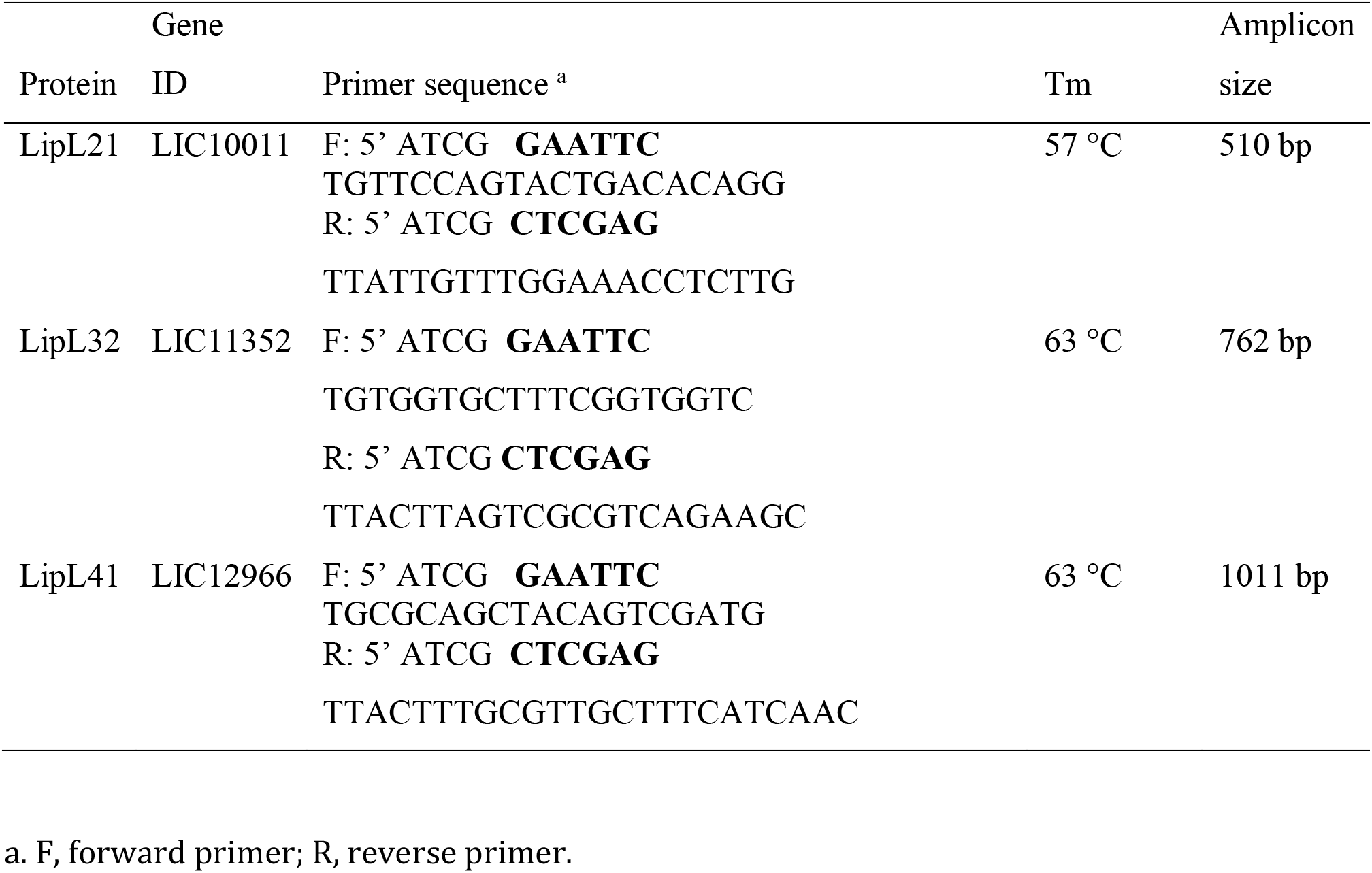
Oligonucleotides primers specific for *lipl21, lipl32* and *lipl41* genes

### Preparation of peripheral blood mononuclear cells (PBMCs)

Blood was collected by jugular vein puncture of cattle into heparin. PBMCs were isolated from blood by layering over Ficoll®Paque Plus (GE Healthcare), followed by centrifugation for 45 min at 400 x g. PBMCs were then washed three times and resuspended in complete RPMI 1640 medium (cRPMI) containing 2 mM L-glutamine, 50 mg/ml gentamicin sulfate, 50 μM 2-mecaptoethanol and 10% (v/v) heat-inactivated fetal bovine serum (Hyclone).

### Cell proliferation assay with [3H]-thymidine incorporation

A total of 5×10^6^ cells/mL was cultured in 96-well flat bottom plates in the presence of 5, 1.5, 0.5 or 0.15 μg/mL complex antigen preparations obtained from *Leptospira* (i.e., WCL, secreted proteins or SDS-soluble protein preparations) or in the presence of recombinant proteins (i.e., LipL21, LipL32, LipL41 or LipL46). Medium, concanavalin A (ConA) at a final concentration of 1 μg/mL or LPS from *E. coli* at a final concentration of 5 μg/mL were used as controls. All cultures were performed in quadruplicate and incubated at 37 °C with 5% CO_2_ for 6 days. Then, 0.25 μCi of [3H] thymidine was added to each well and cultures were incubated for 24 h. Cells were harvested onto glass fiber filter paper by using a cell harvester and the incorporation of [3H] thymidine was measured by liquid scintillation counting. Results were measured in counts per minute (cpm) of [3H] thymidine incorporated into DNA. Results are expressed as mean ± SEM of the cpm in quadruplicate cultures.

### Flow cytometry to determine T cell subsets proliferating in response to antigens of *Leptospira*

Cell division in T cell subsets was evaluated by using the eFluor670 dye. PBMC at 2×10^7^ cells/mL were mixed with 5 mM eFluor670 dye in 1x PBS and incubated for 10 min at 37 °C. After incubation, 10 mL ice-cold cRPMI was added to the cell suspension and an incubation of 10 min was performed on ice. Cells were washed twice and reconstituted at a concentration of 5×10^6^ cells/mL, which was aliquoted into a 24-well tissue culture plates. PBMCs were cultured for 7 days in the presence of 0.15 *μ*g/mL antigens from *Leptospira* or 10 *μ*g/mL recombinant proteins LipL21, LipL32 and LipL41. Medium, ConA and sonicated *L. borgpetersenii* serovar Hardjo antigen (Lepto) were used as controls. After 7 days incubation at 37 °C with 5% CO_2_, cells were stained by indirect immunofluorescence and T cell subpopulations were evaluated by using monoclonal antibodies (mAb) for the cell surface markers CD4 (ILA-11), CD8 (ILA-51), *δ* TCR (GB21A), WC1 (CC15 or IL-A29), WC1.1 (BAG25a) and WC1.2 (CACTB32A) purchased from the Washington State University Monoclonal Antibody Center (https://vmp.vetmed.wsu.edu/resources/monoclonal-antibody-center), followed by staining with goat anti-mouse phycoerythrin or fluorescein-conjugated isotype specific secondary antibodies. Cells were washed, fixed with 4% paraformaldehyde and analyzed by flow cytometry. The viable lymphocytes were determined by the forward and side scatter profile and positive cell percentages and cell division were determined.

### Purification of WC1^+^ *γδ* T-cell and Western blotting for detection of leptospiral protein interaction

WC1^+^ *γδ* T cells were purified from PBMC by magnetic bead sorting (MACS; Milteny Biotec). PBMCs were suspended in PBS containing 6% heat inactivated bovine serum, incubated with mAb ILA-29 (a-WC1) for 30 min at 4 °C, and washed with PBS containing 0.5% bovine serum albumin and 2mM EDTA. Goat anti-mouse IgG microbeads were added, and cells were purified over LS-separation columns (Miltenyi). A total of 2×10^5^ WC1^+^ *γδ* T-cells were resuspended with PBS containing protease inhibitors in a final volume 200 *μ*l. Following this, cells were incubated with 10 *μ*g, 5 *μ*g or 1 *μ*g of LipL21 or LipL32 protein for 1 h at 37 °C with 5% CO_2_ with shaking. Reaction mixtures were centrifuged at 400 x g for 10 min, the supernatant was collected, and the resulting pellet was resuspended in cRPMI. Unlabeled cells also were collected and incubated with proteins. SDS-PAGE gel fractionated proteins were transferred onto PVDF membranes and recombinant protein detection was performed by using anti-his antibodies as described above. Cells incubated only with PBS and recombinant protein alone were used as controls.

### Interaction of recombinant proteins to individual WC1 domains

ELISA plates containing 96 wells (Nunc MaxiSorp, Thermo Fischer Scientific) were coated with 1 μg/well of the WC1-3a1, WC1-3b2, WC1-3d6, WC1-3c8 and WC1-3e10 domains and SUMO tag was used as control. After incubation overnight at 4 °C, plates were washed three times with PBS containing 0.05% Tween 20 (PBS-T) and incubated with a blocking solution containing PBST with 5% BSA. Then, 1 μg/well of recombinant proteins LipL21 and LipL32 were added to the plates for 2h at room temperature for allowing interaction with domains. Plates were washed again and polyclonal antibodies against each recombinant protein were added on dilution of 1:800 for LipL21, and 1:1,000 for LipL32. After 1 hr incubation, binding detection was performed using HRP-conjugated goat anti-mouse IgG (1:5,000), followed by the addition of citrate phosphate buffer (150 mM, pH 5.0) containing 1 mg/mL of o-phenylenediamine and 0.03% H_2_O_2_. The reaction was stopped after 15 minutes, by adding 2 M H_2_SO_4_, and the absorbance (OD_492nm_) was measured using a Multiskan-FC microplate reader (Thermo Fisher Scientific, Helsinki, Finland). Ligation of the recombinant proteins was compared to the SUMO control.

### Statistical analysis

Results are expressed as the mean and SEM for replicate test wells. One-way ANOVA multiple comparisons was used to determine the significance of differences between means and p<0.05 was considered statistically significant. Three to two independent experiments were performed for each assay, each one with triplicate or quadruplicate samples.

## Results

### Antigen preparations containing proteins from *L. interrogans* serovar Copenhageni induce γδ T cells proliferation in cattle immunized against leptospirosis

It has been shown that both human and bovine γδ T cells are able to proliferate and produce IFNγ in response to *Leptospira* (Brown et al., 2003; Klimpel et al., 2003; Naiman et al., 2001). However, the leptospiral components involved in this interaction are still unknown. Thus, we first verified the ability of PBMC from leptospira-immunized cattle to proliferate in response to protein antigens obtained directly from *L. interrogans*. To do that, *Leptospira* cultures were extracted for the following: (i) SDS-soluble cell wall proteins, (ii) secreted proteins and (iii) WCL. Estimated protein concentrations for each fraction were 0.089 μg/μl, 0.2 μg/μl and 0.62 μg/μl, respectively, and the protein profiles in SDS-PAGE gradient gels are shown (Fig. 1). To assess their ability to induce recall responses by PBMC a total 5×10^6^ cells/mL were cultivated with 5, 1.5, 0.5 or 0.15μg/mL of antigens for 7 days. Cultures with ConA or whole cell sonicates of *L. borgpetersenii* serovar Hardjo were used as positive controls since they are known to promote cellular proliferation of PBMC from immunized cattle (Naiman et al., 2001). All three antigen preparations described above promoted proliferation above that in cultures with medium alone (Fig. 2). Proliferation increased with decreasing antigen concentrations; in all cases the lowest concentration of antigen preparations induced more proliferation than either of the positive controls although the difference may not have been significant (Fig. 2). Since the bovine PBMCs responded to our antigen preparations as well as they did to the positive control of sonicated leptospires we have used in our previous studies, we set out to characterize the cell subpopulation(s) involved in responses to these antigens. PBMCs from vaccinated animal were cultured with 0.15 μg/mL of the antigen preparations and after 7 days of culture stained with antibodies to the cell surface markers CD4, CD8, δ TCR and WC1 and analyzed by flow cytometry. As previously observed for the positive control of sonicated *L. borgpetersenii*, antigenic preparations prepared in this study also resulted in inducing a high percentage of CD4 T cells in the cultures and a decrease in CD8 T cells below that in medium controls (Fig. 3B). For WC1^+^ *γδ* T cells, when we compare with medium alone, there was not difference in the number of produced cells (Fig. 3A), but their ability of proliferation when evaluated for cell division by eFluor670 loading (Fig. 3B). The response obtained to WCL fraction was already expected since this fraction also consisted of sonicated leptospira, similar to the *L. borgpetersenii* antigen used historically in previous studies (Blumerman et al., 2007; Naiman et al., 2001; Naiman et al., 2002). In the case of secreted proteins and SDS-soluble protein preparations, this type of response was demonstrated for the first time, suggesting that WC1^+^ *γδ* T cells respond in recall responses to protein antigens.

**Fig. 1.**
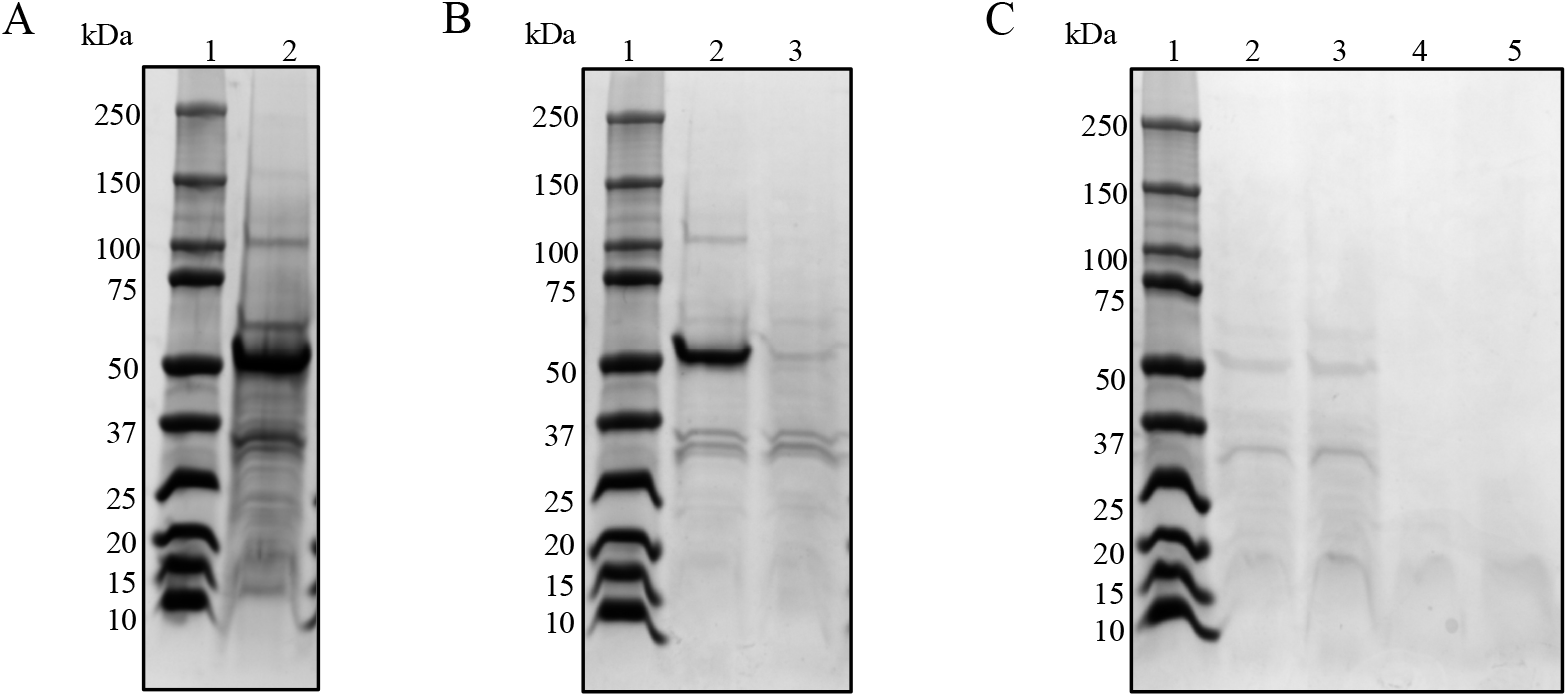
SDS-PAGE gel profile of purified antigens from *L. interrogans* serovar Copenhageni. Gels were stained with Commassie Blue. Lane 1: molecular mass protein markers in all panels. (A) Lane 2, WCL. (B) Lane 2, secreted proteins after 4 h incubation; lane 3, secreted proteins after overnight incubation. (C) Lanes 2 and 3, SDS-soluble proteins after the first extraction; lanes 4 and 5, SDS-soluble proteins after the second extraction. Two lanes were run for each sample type.

**Fig. 2.**
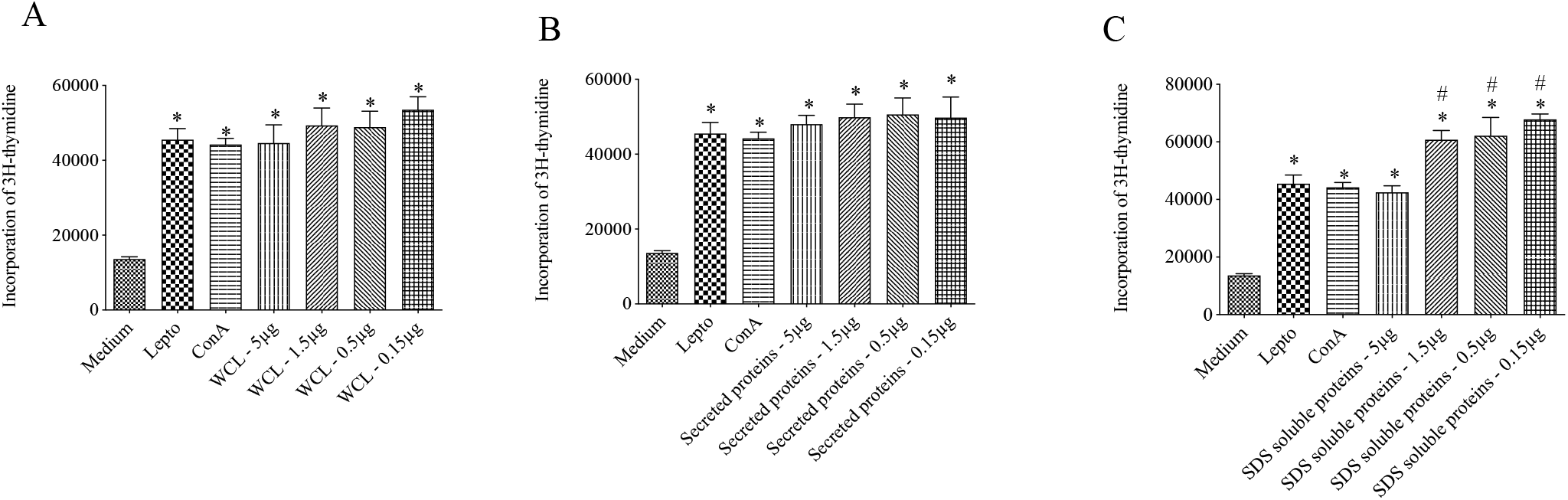
Proliferative responses of PBMC from vaccinated cattle induced by purified antigens from *L. interrogans* serovar Copenhageni. Following culture with several concentrations of (A) WCL fractions, (B) secreted proteins or (C) SDS-soluble proteins proliferative responses were evaluated by incorporation of [3H]-thymidine. Medium, *Leptospira borgpetersenii* sonicate (Lepto) and ConA were used as controls to induce proliferation. Results are expressed as mean ± SEM; asterisks indicate *p* ≤0.05 for proliferative response that were significantly different from those in medium controls and # indicate *p* ≤0.05 for proliferative response that were significantly different from those in Lepto, ConA and the higher concentration of antigen preparation.

**Fig. 3.**
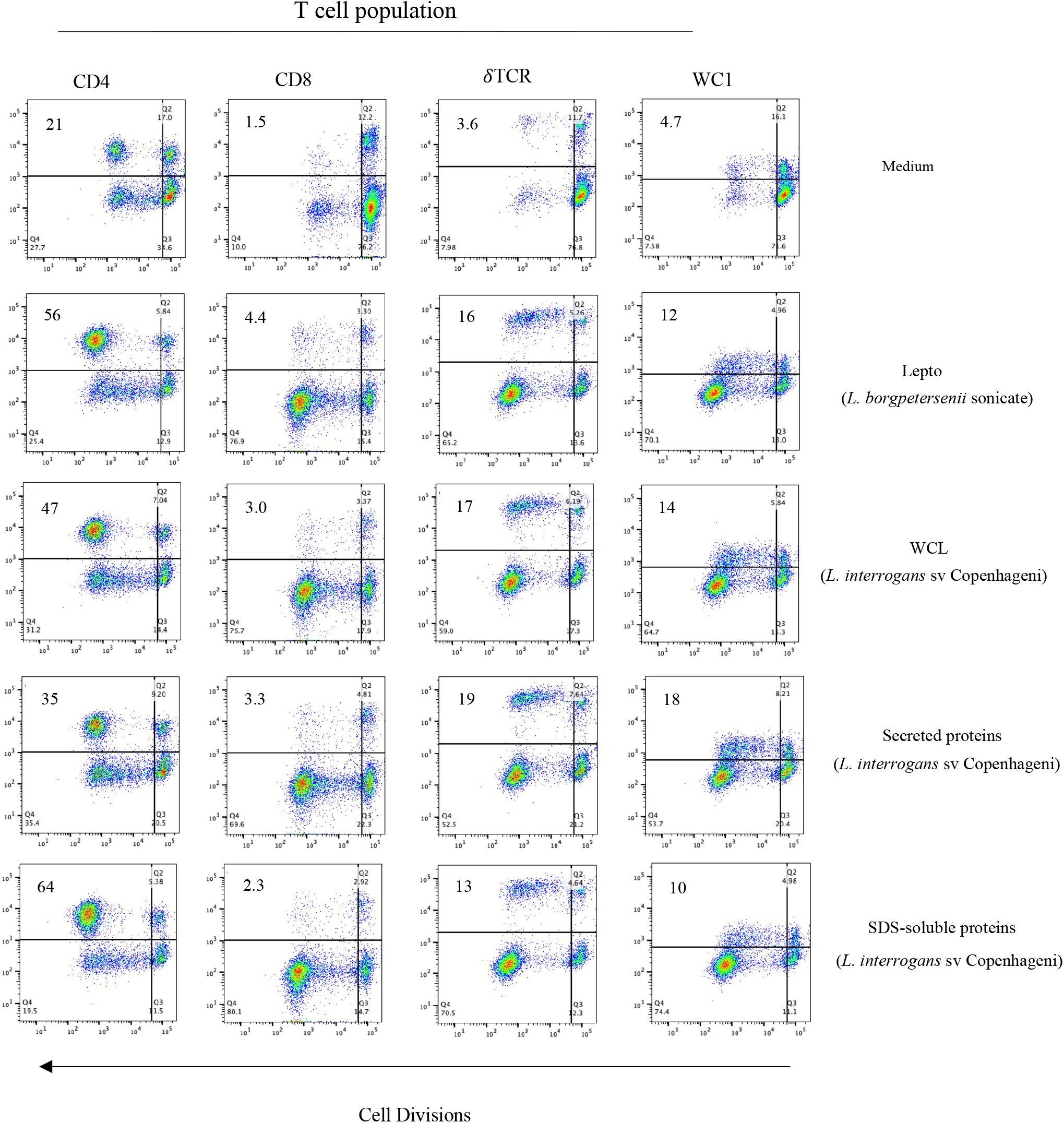
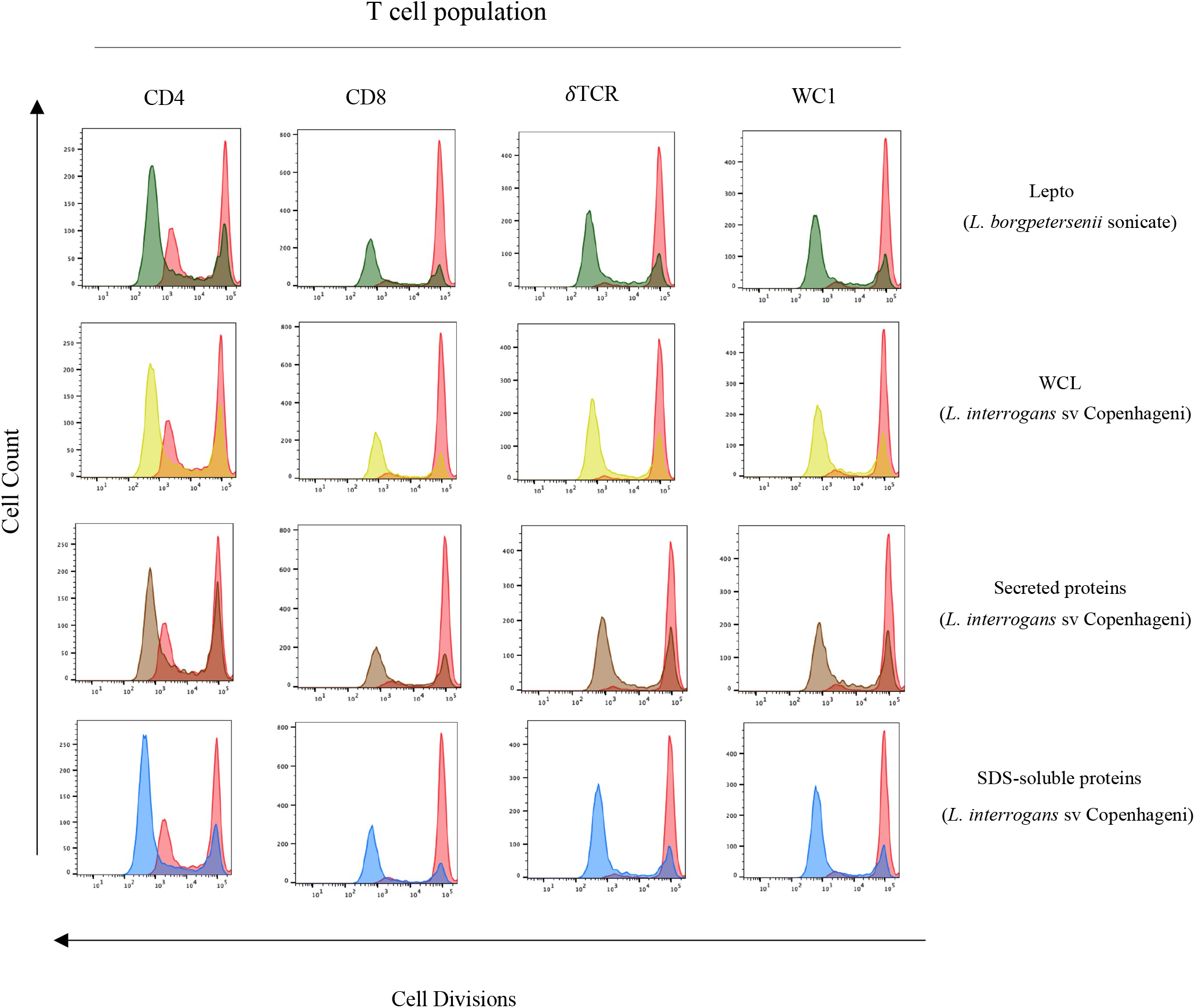
Proliferation of CD4, CD8, *γ*δ T cells and WC1^+^ *γ*δ T cells in response to complex leptospiral antigen preparations measured by eFluor 670 labeling. (A) PBMC from vaccinated cattle were cultured for 7 days with culture additives shown, then stained with the cell surface markers CD4, CD8, *γδ* TCR and WC1 and analyzed by flow cytometry. Sonicated *L. borgpetersenii*, medium and ConA were used as controls. (B) Histograms show the proliferative response profile among the different groups shown in A. Results are representative of three independent experiments.

### Recombinant proteins induce less of a proliferative response compared to extracted antigens, but an expressive response of γδ T cells

LipL21, LipL32 and LipL41 are leptospiral abundant outer membrane lipoproteins, antigenically conserved among pathogenic species and expressed during infection of mammalian hosts (Haake et al., 1999; Haake and Zückert, 2015; Malmström et al., 2009). Those proteins, that are located on the surface, could act as PAMPs and be recognized by specific PRR. Thus, to evaluate the role of recombinant proteins in activation of WC1^+^ *γδ* T cells, the *lipl21, lipl32 and lipl41* genes from *L. interrogans* serovar Copenhageni strain M20 were amplified and cloned into the pET28a vector and used to transform the *E. coli* BL21 (DE3) Star expression strain. The recombinant proteins were expressed with a 6-his tag and purified by immobilized-metal affinity chromatography. SDS-PAGE gel and Western blotting showed that LipL21 and LipL32 proteins were expressed both in soluble and insoluble form (Fig. 4A and 4B, respectively), while LipL41 appeared exclusively in insoluble form (Fig. 4C). All proteins were purified from the insoluble fraction and they were refolded by dilution. After dialysis, an aliquot of each purified protein was analyzed and showed expected sizes of 21, 32 and 41 kDa, respectively (Fig. 4D).

**Fig. 4.**
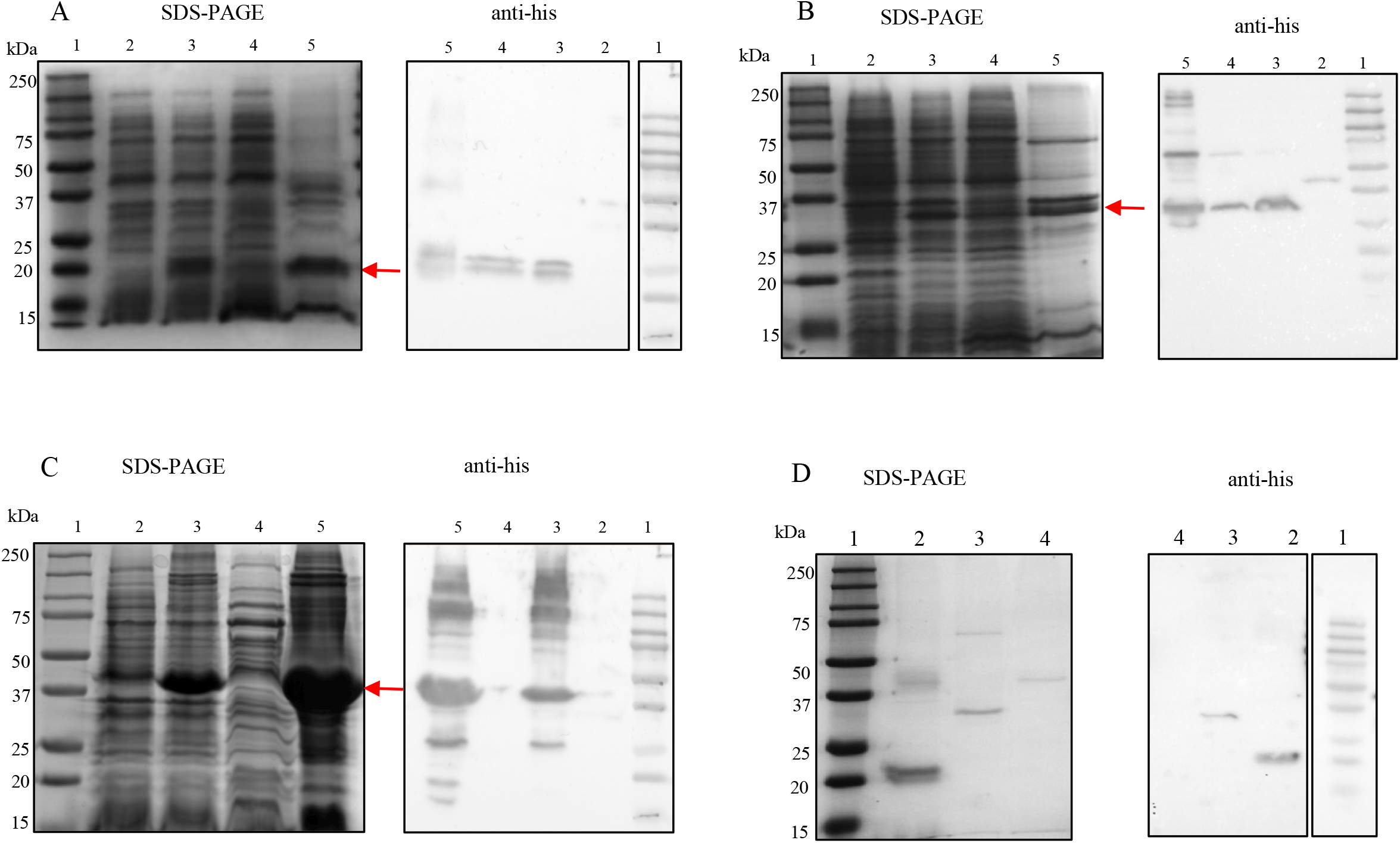
SDS-PAGE gels and Western blots of the expressed and purified recombinant proteins. Expression of recombinant proteins LipL21 (A), LipL32 (B) and LipL41 (C) was done in *E. coli* BL21 DE3 strains and analyzed by SDS-PAGE (left-hand panels) and Western blotting probed with anti-His tag antibodies (right-hand panels). In panels A – C lanes are: molecular mass protein marker (1); non-induced total bacterial extract (2); induced total bacterial extract (3); soluble fraction (4); insoluble fraction (5). The recombinant proteins were purified by affinity chromatography and an aliquot of each protein was analyzed by SDS-PAGE and Western blotting (D). Lanes: molecular mass protein marker (1); LipL21 (2); LipL32 (3); LipL41 (4).

Recombinant proteins were then evaluated for their ability to promote proliferation of PBMCs. Similarly, to extracted antigen preparations, cells were cultivated with several concentrations of recombinant proteins. In contrast to antigens purified directly from leptospires, recombinant proteins induced a reduced cell proliferation (Fig. 5). Only LipL21 at the higher concentration promoted proliferation that was statistically significant different from responses with culture medium alone. However, when the T cell subpopulation was investigated, we observed that all proteins exhibited an increase in the percentage of WC1^+^ *γδ* T cells (Fig. 6). Interestingly, the CD8 T cells profile was different than that observed with extracted antigens since recombinant proteins showed a high tendency to stimulate this cell population. For CD4 T cells, there was no difference in the number of cells in cultures with LipL32 and medium alone, while for the LipL21 and LipL41 proteins the number of cells was less. When cell division was assessed, it was found that LipL32 caused responses by CD4^+^ T cells as well as by TCRδ^+^ cells and particularly by the WC1^+^ γδ T cell subpopulation but not CD8^+^ T cells (Fig. 6). Also, LipL21 stimulated substantial proliferation by the TCRδ^+^ cells as well as some proliferation by the WC1^+^ γδ T cells within that population. Although the LipL41 protein resulted in a high percentage of WC1^+^ *γδ* T cells compared to that in medium alone, when the ability to proliferate was assessed by eFluor670 staining only a very discrete proliferative response was observed.

**Fig. 5.**
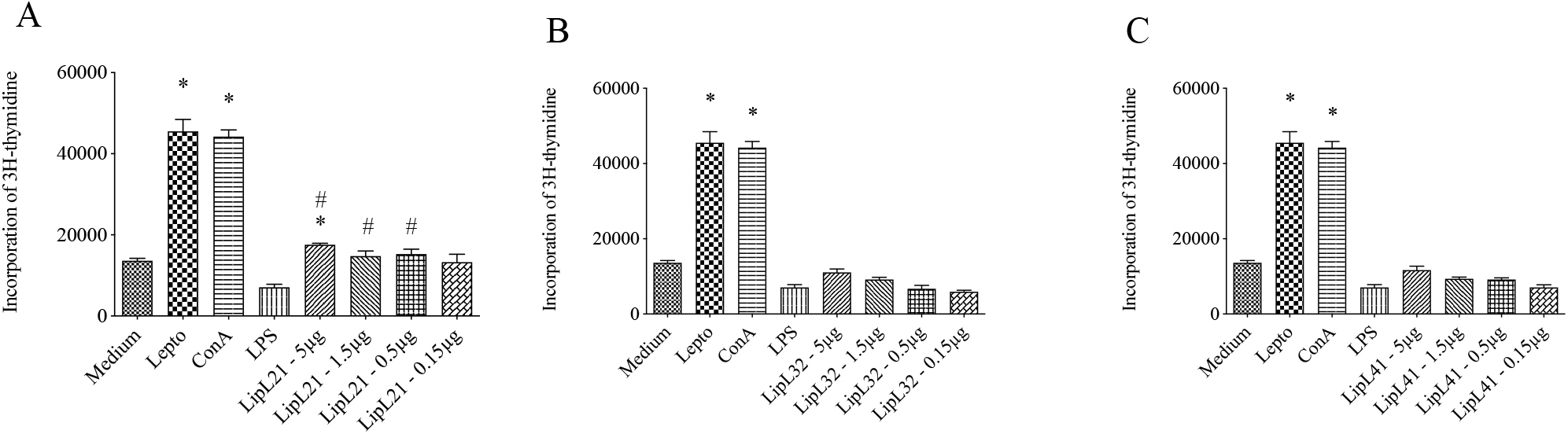
Proliferative response of PBMC induced by recombinant proteins from *L. interrogans* serovar Copenhageni. PBMC were from vaccinated cattle and incubated with several concentrations of recombinant proteins LipL21 (A), LipL32 (B) and LipL41 (C). The proliferative response was evaluated by incorporation of [3H]-thymidine. Medium, Lepto, LPS and ConA were used as control. Results are expressed as mean ± SEM; asterisks indicate a *p* ≤0.05 for proliferative responses that were significantly different to those in medium controls and # indicate *p* ≤0.05 for proliferative response that were significantly different from those in LPS.

**Fig. 6.**
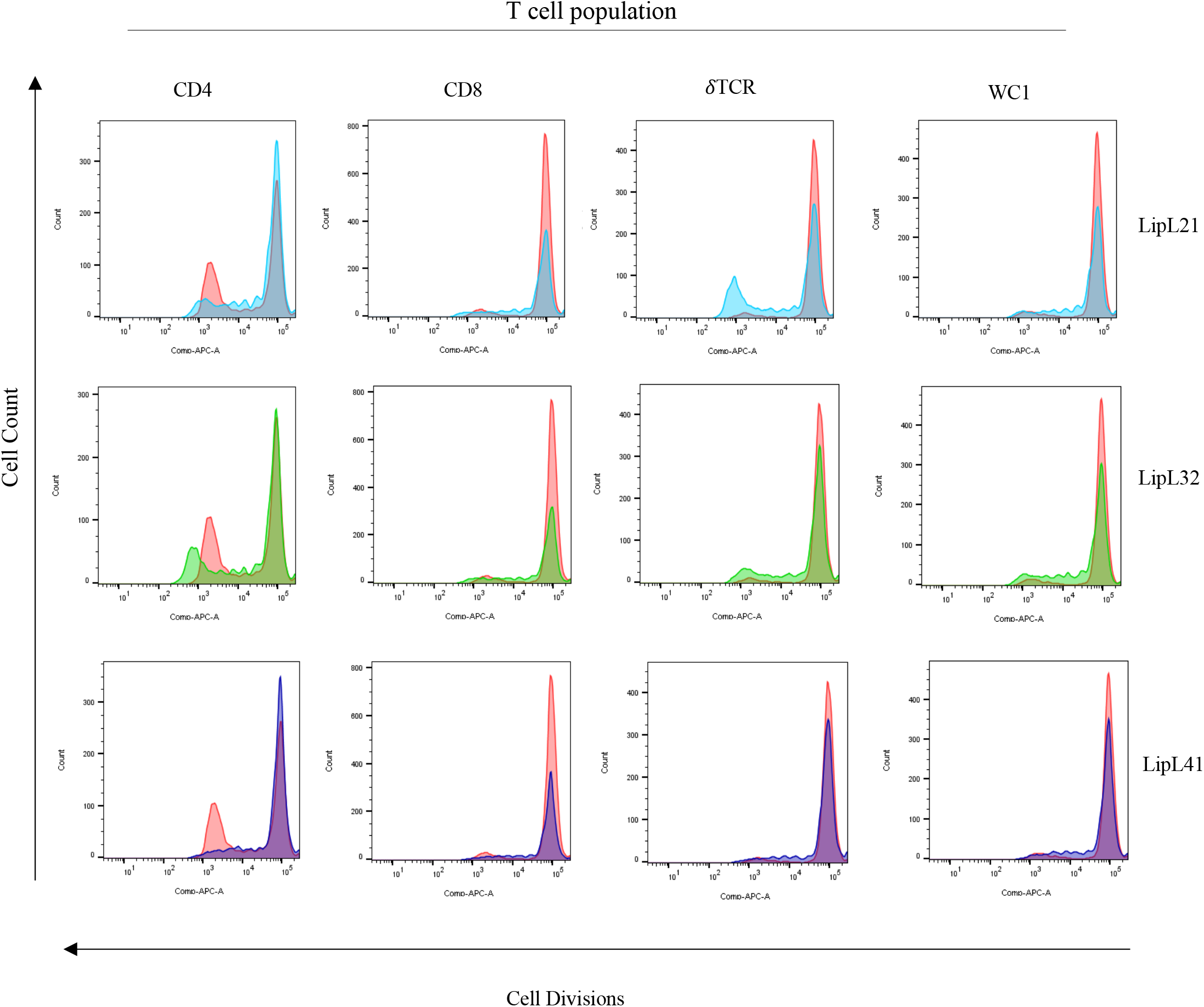
LipL21 and LipL32 show increased ability to induce cell proliferation. PBMC from vaccinated cattle labeled with eFluor 670 were cultured for 7 days in the presence of 10 *μ*g/mL recombinant proteins LipL21, LipL32 and LipL41, then evaluated for expression of CD4, CD8, γ*δ* TCR, or WC1 and analyzed by flow cytometry. Histograms show the proliferative response profile for medium (red line) and recombinant protein indicated (blue line for LipL21, green line for LipL32 and purple line for LipL41). Results are representative of two to three independent experiments.

### WC1.1^+^ *γδ* T-cell subpopulations are involved in response to recombinant proteins

Although the potential of recombinant proteins to promote cell proliferation was reduced compared to the crude antigen extracts, when we explored the T cell subpopulation responses all recombinant proteins induced proliferation by WC1^+^ *γ*δ T cells. Thus, we set out to verify which WC1^+^ *γ*δ T cells subpopulations are involved in response to protein antigens, since WC1.1 and WC1.2 are expressed on mostly non-overlapping *γ*δ T cell populations and it has been reported by us previously that WC1.1^+^ cells are largely the cells that proliferate and produce IFN-*γ* in leptospira antigen-stimulated cultures (Naiman et al., 2001; Rogers et al., 2005a). As shown in Fig. 7, the percentage of both WC1.1^+^ and WC1.2^+^ *γδ* T-cell proliferating to the recombinant proteins was higher than in cultures of medium alone. However, the WC1.1^+^ *γδ* T-cell response was greater for all three proteins although LipL41 showed a slightly less stimulation than LipL21 and LipL32. Unlike WC1.1^+^ cells, the percentage of WC1.2^+^ positive cells were also higher in response to recombinant proteins, albeit, less than that by WC1.1^+^ cells. The observation that WC1.2^+^ cells respond to *Leptospira* proteins is interesting since previously we showed that sonicated leptospires generally promoted a response by WC1.1^+^ *γδ* T-cell (Rogers et al., 2005a).

**Fig. 7.**
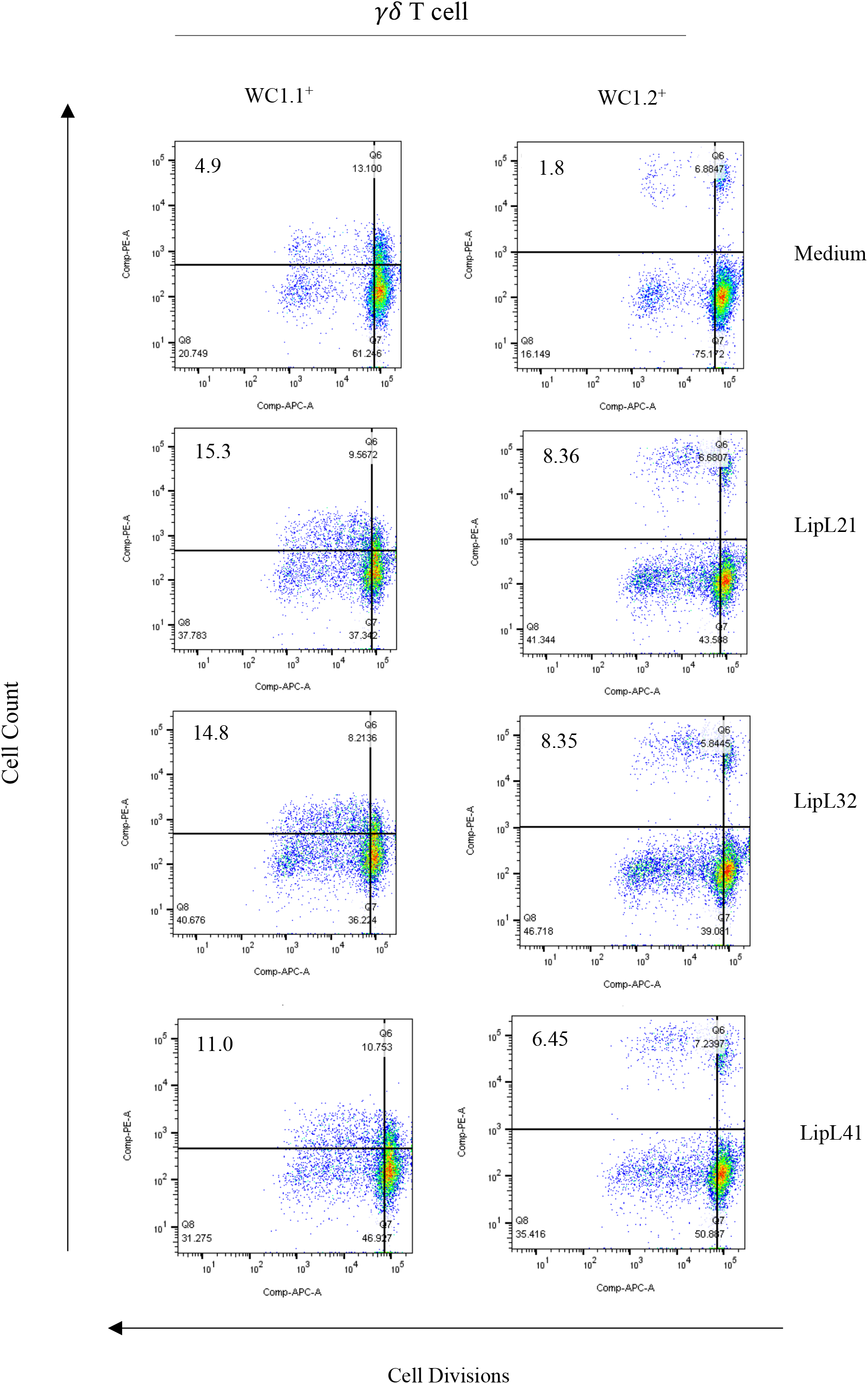
WC1.1^+^ *γδ* T-cell subpopulations respond to protein antigens from *Leptospira*. PBMC from vaccinated cattle labeled with eFluor 670 were cultured for 7 days in the presence of 10 *μ*g/mL recombinant proteins LipL21, LipL32 or LipL41, and stained with anti-WC1.1 (mAb BAG25A) or anti-WC1.2 (mAb CACTB32). The expression of WC1.1 and WC1.2 was analyzed by flow cytometry. The left panels show the percentage and proliferation of WC1.1^+^ *γδ* T-cell in response to antigens while in the right panels is shown response obtained for WC1.2^+^ *γδ* T-cell. Medium only cultures were used as controls. Results are representative of two or three independent experiments.

### The LipL21 and LipL32 recombinant proteins interact directly with WC1^+^ *γδ* T-cell

Aiming to verify if LipL21 and LipL32 proteins interact directly with WC1^+^ *γδ* T cells, we purified WC1^+^ *γδ* T cells and incubated a total 2×10^5^ cells/mL with 10 *μ*g, 5 *μ*g or 2.5 *μ*g of each recombinant protein for 1 h at 37 °C. Unlabeled cells, collected after application of total cells, were also incubated with the recombinant proteins under the same conditions. Moreover, cells incubated with PBS were used as a control. After 1 hr incubation, cells were centrifuged gently, following which the pellet and supernatant were harvested for analysis. As shown in Fig. 8, LipL21 and LipL32 were detected on cell pellet with the three of the conditions used. Excess of proteins were detected on supernatant only in aliquots that were more concentrated. Both LipL21 and LipL32 were detected in a multimeric form, that can have been caused by the lack of a reducing agent in the SDS buffer. As cells incubated only with PBS did not show such bands, unspecific reaction caused by antibodies have been dismissed (Fig. 8 - lane 2 for pellet and lane 6 for supernatant). The band of 20 kDa for LipL21 protein was not detected, perhaps because the low concentration used was not enough for detection. In Fig. 8 A, some interaction was detected in unlabeled cells, but since we used the same protein concentration we can suppose that these interactions were stronger with WC1^+^ *γδ* T cells. Taken together, this is the first study that identifies *Leptospira* recombinant proteins as ligands of WC1^+^ *γδ* T cells.

**Fig. 8.**
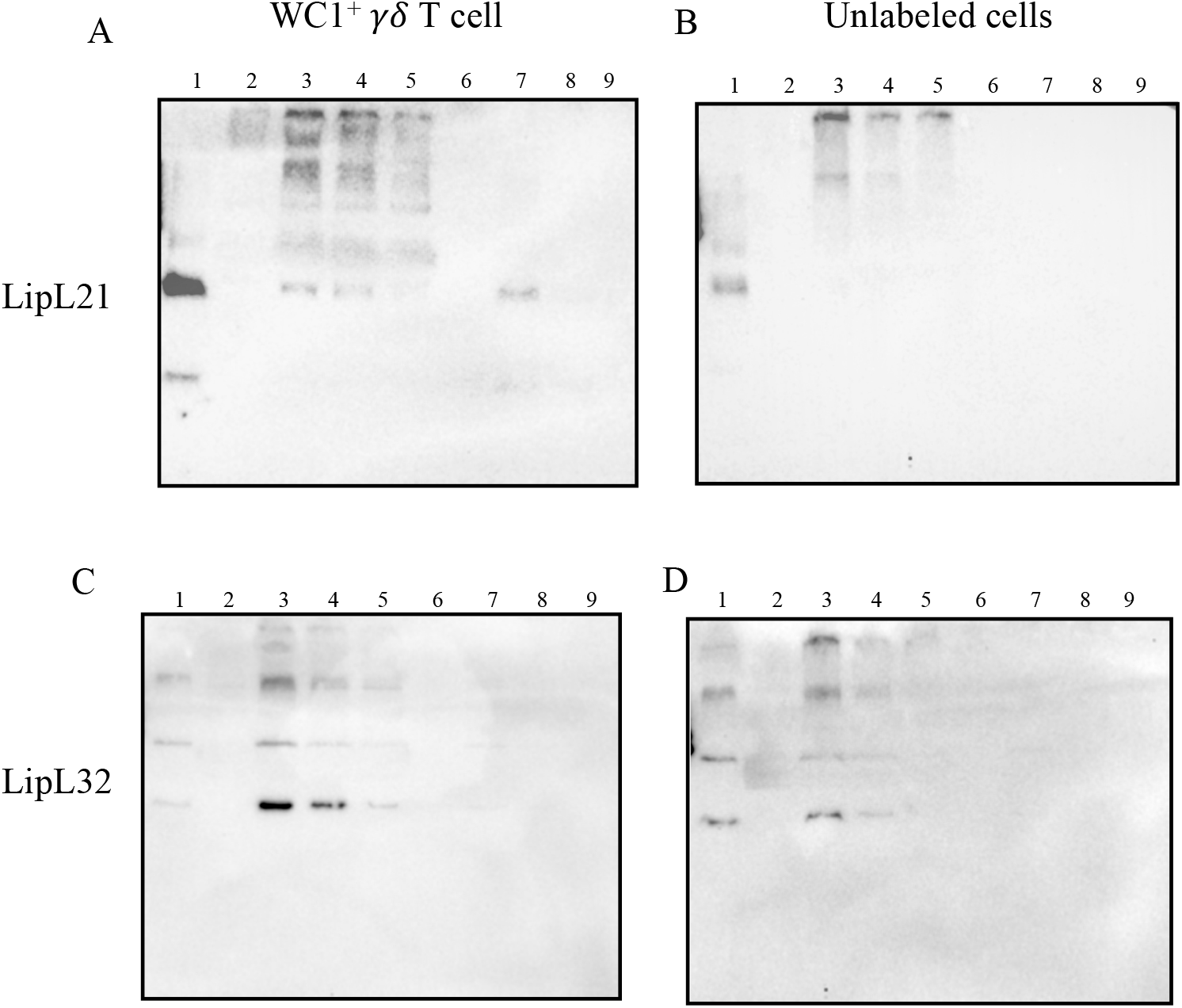
WC1^+^ *γδ* T cells from *Leptospira* vaccinated cattle interact directly with the recombinant proteins LipL21 and LipL32. WC1^+^ *γδ* T cells purified from PBMC by magnetic bead sorting after staining with mAb ILA-29 were incubated with several concentrations of LipL21 and LipL32 for 1 h at 37 °C. Cell suspension containing unlabeled cells also (IL-A29 negative cells) was collected and incubated with the proteins using the same condition. Pellets and supernatants of both preparations were recovered, loaded onto SDS-PAGE for protein fractionation and transferred onto a PVDF membrane. Detection of interaction was performed by using anti-his antibodies and HRP-conjugated anti-IgG. Left-hand panels show the interaction of purified WC1^+^ *γδ* T-cell with LipL21 and LipL32 while the righthand panels show the unlabeled cells (IL-A29^-^) with LipL21 and LipL32 interaction. Lane1, recombinant protein alone; lane 2, cell pellet resulting from incubation with PBS. The cell pellet resulting from incubation with lane 3, 10 *μ*g recombinant protein, lane 4, 5 *μ*g recombinant protein, or lane 5, 2.5 *μ*g recombinant protein. Supernatant resulting of incubation with lane 6, PBS lane 7, 10 *μ*g recombinant protein, lane 8, 5 *μ*g recombinant protein, or lane 9, 2.5 *μ*g recombinant protein.

### LipL21 and LipL32 are potential WC1-3 SRCR domains ligands

WC1 proteins are organized in the SRCR domain pattern of a1-[b2-c3-d4-e5-d6]-[b7-c8-d9-e10-d’11]. Previously, five SRCR domains of WC1-3 were shown by us to interact with vaccine preparations of *L. borgpetersenii* serovar Hardjo-bovis and *L. interrogans* serovar Hardjo-bovis (Hsu et al., 2015). To examine whether the proteins LipL21 and LipL32 contribute to that binding to specific SRCR domains, we cloned, expressed, and purified the SRCR domains WC1-3a1, WC1-3b2, WC1-3d6, WC1-3c8 e WC1-3e10 and evaluated their interaction by ELISA. Expressed SRCR domains were immobilized on high protein-binding capacity plates and LipL21 and LipL32 attachment was evaluated by using polyclonal antibodies against the recombinant proteins. As demonstrated in Fig. 9A (LipL21) and Fig. 9B (LipL32), both proteins exhibited the ability to interact with all SRCR domains tested when compared to binding with SUMO protein, used as negative control (Fig. 9). These observations were confirmed by Western blotting analyses (data not shown), suggesting that the proteins LipL21 and LipL32, which are present in the outer membrane of pathogenic *Leptospira* spp., may contribute to activation of WC1^+^ *γδ* T cells by directly binding to the extracellular WC1 SRCR domains.

**Fig. 9.**
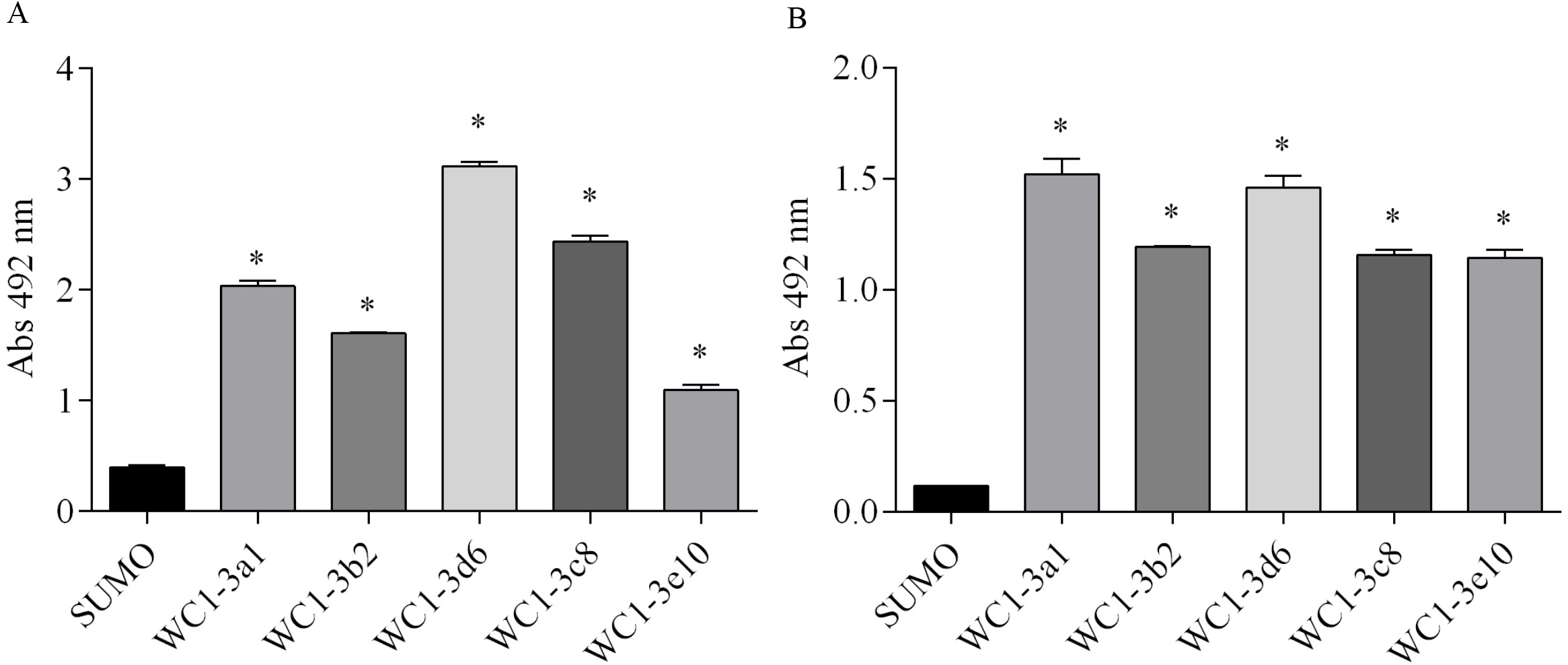
WC1-3’s SRCR domains bind to the recombinant proteins LipL21 and LipL32. ELISA plates were coated with recombinant proteins representing SRCR domains WC1-3a1, WC1-3b2, WC1-3d6, WC1-3c8, WC1-3e10 or with the control protein SUMO. The interaction of recombinant protein LipL21 (A) and LipL32 (B) was detected by addition of anti-recombinant polyclonal murine antibodies. Bars represent the mean ± SEM absorbance at 492 nm of three replicates; asterisks indicate *p* ≤0.05 compared to the binding of recombinant protein to SUMO. Results are representative of three independent experiments.

## Discussion

It was believed that protective immunity to leptospirosis was exclusively humoral since the presence of high titers of anti-LPS antibody specific to specific serovars was able to promote protection against the disease (Faine et al., 1999). However, in 2001 was demonstrated that a vaccine against *L. borgpetersenii* serovar hardjo induced a very strong Th1-type immune response in vaccinated cattle (Naiman et al., 2001) while antibody-inducing vaccines did not protect against this serovar (Bolin et al., 1989). Since then, this type of response has been studied and the involvement of WC1^+^ *γδ* T and CD4 T cells has been shown (Naiman et al., 2001; Naiman et al., 2002). The leptospiral antigens responsible by activation of Th1-type immune response had not been found and it was unclear whether these antigens were protein or non-proteins. This was based on the fact that previous work had shown that treatment of *Leptospira* spp. both with proteinase K and polymyxin B did not prevented the binding of leptospires to the WC1 domains, suggesting that these antigens were unlikely to be either protein or LPS (Hsu et al., 2015) however there were shortcomings with these experiments. Here we showed that protein preparations obtained directly from *L. interrogans* serovar Copenhageni were able to stimulate *γδ* and CD4 T cell proliferation similarly to that induced by whole bacterial cell sonicate. Moreover, two outer membrane proteins able to interact directly with WC1^+^ *γδ* T cells were identified, i.e., LipL21 and LipL32, both of which are major outer membrane proteins of *Leptospira* spp. Although several studies have used proteinase K as a method to examine the surface exposure of proteins in many pathogens (Bunikis and Barbour, 1999; Sabarth et al., 2002; Thomas et al., 1992), some studies performed with surface-exposed leptospiral proteins have shown insensibility to proteinase K cleavage (Cullen et al., 2005; Koizumi and Watanabe, 2003; Pinne and Haake, 2009). In the study presented here, we observed that both proteins when subjected to proteinase K cleavage in their recombinant form were susceptible to cleavage, but when investigated in leptospiral cells both proteins showed resistance to digestion with proteinase K (unpublished data). These data suggest that LipL21 and LipL32 are not accessible to proteinase K cleavage and possibly this inaccessibility is associated to with the steric hindrance caused by LPS structures present on the cell surface.

*γδ* T cells has been described as recognizing a range of antigens including proteins, carbohydrates, lipids and nucleic acids. These molecules could act as potential ligands or at least be involved in the recognition by and activation of these cells (Deseke and Prinz, 2020). In response to *Mycobacterium, γδ* T cells recognize both protein and nonprotein antigens (McGill et al., 2014; Smyth et al., 2001; Vesosky et al., 2004; Welsh et al., 2002). Similarly, to *L. borgpetersenii* both CD4 T cells and WC1^+^ *γδ* T cells from *M. bovis*-infected cattle have been shown to be highly responsive to *M. bovis* sonicated extracts (Smyth et al., 2001). Thus, the response obtained in our study to the WCL fraction was not unexpected, since this fraction consisted in sonicated leptospires. Interestingly, Welsh and collaborators showed that after enzymatic digestion of the proteins present in *M. bovis* sonicated extract the ability to stimulate CD4 T cell responses was abolished and that although this treatment resulted in reduction of WC1^+^ *γδ* T cells proliferation it did not completely eliminate it; this showed that two or more molecules are involved in activation these cells. One of the main protein antigens responsible for activation of CD4 T cells and WC1^+^ *γδ* T cells by *Mycobacterium* is the 6-kDa early secretory antigenic target (ESAT-6) and antigen 85, both isolated from culture filtrate (Rhodes et al., 2000; Rhodes et al., 2001). In our study, the leptospiral protein preparations obtained from bacterial cultures also induced a strong proliferative response by CD4 T and WC1^+^ *γδ* T cells. However, the same potential was not observed when recombinant proteins were evaluated. Similarly, when native and recombinant ESAT-6 preparations were used *in vitro* to stimulate splenic lymphocytes, the native preparation was more active than the recombinant forms in terms of their ability to stimulate IFN-*γ* production. Interestingly, the native preparation induced levels of IFN-*γ* very close to those found by total culture filtrate (Sørensen et al., 1995). We are not sure whether like ESAT-6 that the LipL21 and LipL32 proteins are present in abundance in the culture preparations, but, regardless, our data suggest that the presence of additional or accessory molecules may be the key to activating a potent immune response.

The hypothesis that WC1^+^ *γδ* T cell activation is mediated by co-ligation of *γδ* TCR and WC1 by a pathogen is very well supported by previous works, in which the SRCR domains binding to *Leptospira* were identified (Baldwin et al., 2020; Hsu et al., 2015). Since WC1 transcripts undergo extensive alternative splicing (Herzig and Baldwin, 2009), it is believed that any change in the number of SRCR domains in the extracellular region of WC1 molecules after splicing could affect avidity for the bacteria or ligand and thus reduce WC1/TCR co-ligation. Perhaps the use of a single recombinant protein to stimulate PBMCs from cattle may have been insufficient to promote a high affinity interaction and/or cross-linking of receptors to result in a full activation of the WC1^+^ *γδ* T cell. It could explain the reason LipL21 and LipL32 promoted a less potent response, since we have demonstrated that both LipL21 and LipL32 interact directly with WC1^+^ *γδ* T cells. That is, LipL21 and LipL32 interacted with the five WC1 SRCR domains previously identified as responsible for binding *Leptospira* (Hsu et al., 2015). In that *in vitro* experimental set-up some important antigen epitopes could have been hidden; alternatively as suggested above full activation of WC1^+^ *γδ* T cells may require the use of two or more proteins. We still need to know whether the interaction of LipL21 and LipL32 with WC1^+^ *γδ* T cells also requires binding to the γδTCR thereby engaging both WC1 and TCR. It is anticipated that understanding these receptor interactions will be important to fully understand the ability of these proteins to prime WC1^+^ *γδ* T cells and will contribute to development of new vaccine strategies.

*Leptospira*-responsive WC1^+^ *γδ* T cells has been defined serologically as the WC1.1^+^subpopulation by mAbs that react with WC1 SRCR domains, while the WC1.2^+^ subset has been involved in the recognition of other pathogens (Lahmers et al., 2005; Rogers et al., 2005a; Rogers et al., 2005b). Normally, WC1.1-type and WC1.2-type molecules are not expressed simultaneously by the same cell, suggesting that expression of WC1 family members is directed in a way that confers T cell specificity for the antigen (Chen et al., 2012). However, more recently it was observed that some *γδ* T cell clones express WC1 genes representative of both subsets (Damani-Yokota et al., 2018). We have shown in this work that both WC1.1^+^ and WC1.2^+^ *γδ* T-cell subsets were involved in responses to the *Leptospira* protein antigens, although a greater response by WC1.1^+^ *γδ* T-cell was observed. As some WC1.2^+^ *γδ* T-cell clones produce transcripts coding for WC1 molecules that have a binding domain for *Leptospira* (Damani-Yokota et al., 2018) it is possible that those domains have been responsible for recognition of the proteins LipL21 and LipL32, corroborating the studies that showed the involvement of a small WC1.2^+^ *γδ* T-cell population in responses to *Leptospira* (Rogers et al., 2005b). Price and collaborators (Price et al., 2010), evaluating the role of WC1^+^ *γδ* T-cells in response an intranasal mycobacterial vaccine in calves, also demonstrated a predominant involvement of WC1.1^+^ cells in the lungs of these animals. Nevertheless, another study has shown the presence of both *γδ* T-cell subsets in response to virulent *M. bovis* infection. That is, curiously and in contrast to the results of Price and collaborators with the vaccine strain, an accumulation of WC1.2^+^ *γδ* T-cells was observed in the lungs of these cattle (McGill et al., 2014). While plausible, it is still not clear whether the fact that one of these studies evaluated the immune response after BCG vaccination while the other used a virulent *M. bovis* strain infection influenced the results. In any case, the responses involving the WC1.1 and WC1.2 *γδ* T-cell subsets seems to be a quite complex.

In conclusion, we demonstrated for the first time the involvement the leptospiral protein antigens in the activation of WC1^+^ γδ T cells and identified two leptospiral outer membrane proteins able to interact directly with WC1^+^ γδ T cells. As WC1 has been considered the main element regarding pathogen recognition, we suggest that the protein pools extracted from *Leptospira* promoted the co-ligation of WC1 with the γδ TCR resulting in a higher affinity interaction and, consequently, in a potent immune response. Since we used, as the model, cattle immunized with inactivated leptospires perhaps for that reason a robust response has not been observed after using recombinant proteins. But since they were able to interact with WC1^+^ γδ T cells, we suggest that both LipL21 and LipL32 could be used as ligands to engage γδ T cells in prophylactic immune responses. Certainly, more studies are necessary to verify whether crosslinking of WC1 with the γδ TCR by these proteins will result in the production of proinflammatory cytokines and consequently activation of conventional adaptative immune responses. In general, the understanding of antigenic targets that contribute to stimulation of non-conventional lymphocytes will significantly improve the potential for development of new vaccine strategies.

## Acknowledgments

The research was supported by AFRI Grant #2016-09379 from the USDA-National Institute of Food and Agriculture to C. Baldwin and Hatch funding from the Umass Center for Agriculture, Food and the Environment under project #2016-67015-24913 to C. Baldwin and J. Telfer. AFT was supported by a fellowship from FAPESP (2019/05466-4). The funders had no role in study design, data collection and analysis, decision to publish, or preparation of the manuscript.

AFT contributed to the design of experiments, acquisition, analysis, interpretation of the data and drafting the manuscript. AG, AY and EB contributed to the design of experiments of cell proliferation by using [3H]-thymidine incorporation and flow cytometry, acquisition and analysis of these data. CB, JT and ALTO contributed to the conception and design of the research, interpretation and drafting the manuscript. All authors reviewed the results and approved the final version of the manuscript.

